# A noble TGFβ biogenesis inhibitor exhibits both potent anti-fibrotic and anti-inflammatory capabilities

**DOI:** 10.1101/770404

**Authors:** Han-Soo Kim, Moon Kee Meang, Saesbyeol Kim, Ji Yong Lee, Baik L. Seong, Ik-Hwan Kim, Byung-Soo Youn

## Abstract

Idiopathy pulmonary fibrosis (IPF) is an intractable and fatal human disorder. Our previous study showed that eupatilin exerted a potent anti-fibrotic effect on both in *vitro* fibrogenesis and bleomycin-induced lung fibrosis model (BLM). Subsequently, an analog called ONG41008 had been identified as a more potent anti-fibrotic than eupatilin and also showed a potent anti-inflammatory capability. Orally administered ONG41008 significantly improved onset of BLM in both prophylactic and therapeutic model and its therapeutic efficacy was similarly compared to or better than pirfenidone by measuring production of collagen and hydroxyproline. Staining collagen or αSMA corroborated these results.

As *in vitro* fibrogenesis models, DHLF (Diseased Human Lung Fibroblasts from IPF patients) and HSC (hepatic stellate cells) were used for direct effects of ONG41008 on pivotal cellular and molecular functions associated with pathogenic myofibroblasts; ONG41008 dismantled latent TGFb complex (LTC), generating inactive forms of TGFβ, likely limiting TGFβ to TGFβ receptor via depolymerization of F-actin and this blunted SMAD2/SMAD3 phosphorylation, thereby reprogramming EMT. A set of cell imaging studies and transcriptomic analysis were conducted to explore how ONG41008 elicited both anti-fibrotic and anti-inflammatory capabilities. Elastin (ELN) seemed to be a pioneering pharmacodynamic marker. It was also found that NOX4 played an important role in anti-fibrosis because it was functionally connected to major central nod proteins such as lysyl-oxidase (*LOX*) and numerous collagen family members in an ONG41008-specific fibrogenic interactome. Human *NOX4* was significantly induced by TGFβ and completely knocked down by ONG41008. It has been shown that production of reactive oxygen species (ROS) led to activation of inflammasome. ONG41008 may be likely related to anti-inflammation, leading to a key protective effect on fibrogenesis. Concomitant with downregulation of *NOX4*, expression of macrophages homing chemokines, *CCL2 and CCL7* were significantly attenuated by ONG41008. *In vitro* anti-inflammatory activities of ONG41008 were investigated in RAW264.7 cells, a mouse monocytic cell line stimulated with LPS. ONG41008 substantially attenuated *TNFα*, *CXCL10*, *CCL2* and *CCL7*, which are proinflammatory cytokine and important chemokines influencing T cells or macrophages. TNFα was situated at the central nod in LPS-treated macrophages via an ONG41008-specific interactome analysis.

Taken together, ONG41008 is a TGFβ biogenesis inhibitor, being a potent drug for a broad range of fibrotic diseases and could antagonize inflammatory diseases as well.

## Introduction

Idiopathy pulmonary fibrosis (IPF) is defined a rare disease belonging to interstitial lung diseases (ILD) (1). Its mortality and morbidity are becoming significant such that the median survival of patients with IPF is 3∼5 years after diagnosis and the majority of patients would succumb to death within 5 year of diagnosis (2). Development of both potent anti-fibrotic and anti-inflammatory drugs is of paramount importance for dealing with IPF.

Myofibroblasts (MFB) play a central role in the initiation and perpetuation of fibrosis (3). While identification of the IPF-initiating cells remains to be discovered HSCs are the major cell types contributing to both liver inflammation and liver fibrosis (4). Although fibrogenic signature proteins such as Collagens, CTGF (connective tissue growth factor) or Periostin have been well elucidated, full spectrum of fibrogenic proteins have yet to be discovered. Nevertheless, several anti-fibrotic modalities have continually put on clinical studies (5). Although fibrosis is not defined an immune disorder, inflammatory cells are believed to be responsible for eliciting fibrogenic signaling. Of these cells, macrophages including bone-marrow-derived ones or tissue residential macrophages like alveolar macrophages in the lung may play important roles in establishing early stage inflammation by producing proinflammatory cytokines such as TNFα or IL-1β and probably indirectly affecting TGFβ (6). Therefore, controlling these proinflammatory cytokines or TGFβ may lead to an efficient modality for fibrotic diseases.

Flavones are members of the polyphenol family, a group of over 10,000 compounds have been exclusively found in the plant kingdom (7). In general, these phytochemicals protect plants from radiation damage (8). Due to their anti-oxidant and anti-inflammatory potentials, flavones have long been used to treat inflammatory diseases such as arthritis and asthma (9).

Chromone, 1,4-benzopyrone-4-one, is a central chemical scaffold hereinafter called chromone- scaffold (CS) constituting flavones and isoflavones (10), and the CS derivatives (CSD) are a diverse family based on branching chemical residues coupled to the core CS framework (11). We recently reported that eupatilin from an *Artemisia* species dramatically inhibited LPS-induced osteoclastogenesis via actin depolymerization (12) and downregulation of multiple genes involved in EMT. And eupatilin exhibited powerful anti-fibrogenic potential as well as anti-fibrotic activities in BLM (13). Here, we show that a noble synthetic CSD analog, called ONG41008, was able to reprogram EMT via dismantling LTC, resulting in inactive TGFβ, which significantly ameliorated lung fibrosis in BLM. ONG41008 was able to abrogate production of TNFα and chemokines in activated macrophages, suggesting that ONG41008 is a potent anti-inflammatory drug. ONG41008 may open a door as a powerful new therapeutic modality for treating IPF as well as autoreactive T cells-mediated hyper-inflammation.

## Results

### ONG41008 dismantled latent TGFβ complex (LTC) via depolymerization of F-actin

When DHLF (diseased human lung fibroblasts from IPF patients) or ONGHEPA1 (mouse HSC cells) were stimulated with TGFβ, expression of αSMA (α-smooth muscle actinin) was notable via ICC (immune cell chemistry) but was then substantially mitigated with the treatment of ONG41008 in DHLF (Figure 1A) and in ONGHEA1 (Figure 1B), respectively. Be noted while DHLF resembled pathogenic myofibroblasts proven by an entire transcriptome analysis ONGHEPA1 are hepatic mesenchymal stem cells being trans-differentiated into myofibroblasts in the presence of TGFβ. Its transcriptome was also analyzed. These data are to be seen in the following figure or a supplementary figure. Thus, ONGHEPA1 could be optimal cells for researching on *in vitro* fibrogenesis. And DHLF are relevant cell types for screening anti-fibrotic drugs. A plethora of fibrinogenic markers involving C*OL1α1*, C*OL11α1* or *PERIOSTIN* were induced by TGFβ and were completely knocked down by ONG41008, suggesting that ONG41008 may directly act on pathogenic myofibroblasts or HSC. (Figure 1A and 1B). ONG41008 markedly inhibited phosphorylation of both SMAD2/ SMAD3, suggesting that phosphorylated Smad2 /Smad3 may translocate to the nuclei, activating EMT. Vibrant phosphorylation of ERK with the use of ONG41008 remained to be further discovered (Figure 1C). In our previous study, we showed that eupatilin effectively depolymerized F-actin (12). Accordingly, we hypothesized that F-actin in actin filaments could be also a molecular target for ONG41008. As shown in Figure 1D, actin filaments were apparent in TGFβ- stimulated cells and the severed actin fragments were uniformly distributed in the cytoplasm upon ONG41008 treatment in the presence of TGFβ. While a massive actin depolymerization was observed with the treatment of ONG41008 no such depolymerization was seen in ONGHEAPA1 cells stimulated with pirfenidone or nintedanib, suggesting that the latter two anti-IPF drugs have no effects on F-actin depolymerization. *In vitro* actin polymerization or depolymerization assay was conducted such that ONG41008 was, indeed, an inhibitor of actin polymerization (Figure 1E) as well as an enhancer of actin depolymerization (Figure 1F). It has well been appreciated that osteoclasts exist as cellular fusion states termed multinucleation whereby the O-ring made of F-actin surrounds the fused multinucleated osteoclasts (14). As shown in Figure 1G, ONG41008-treated multinucleated osteoclasts gave rise to massive actin depolymerization. These studies indicate that ONG41008 may be a bona-fide F-actin depolymerizer.

**Figure. 1.**
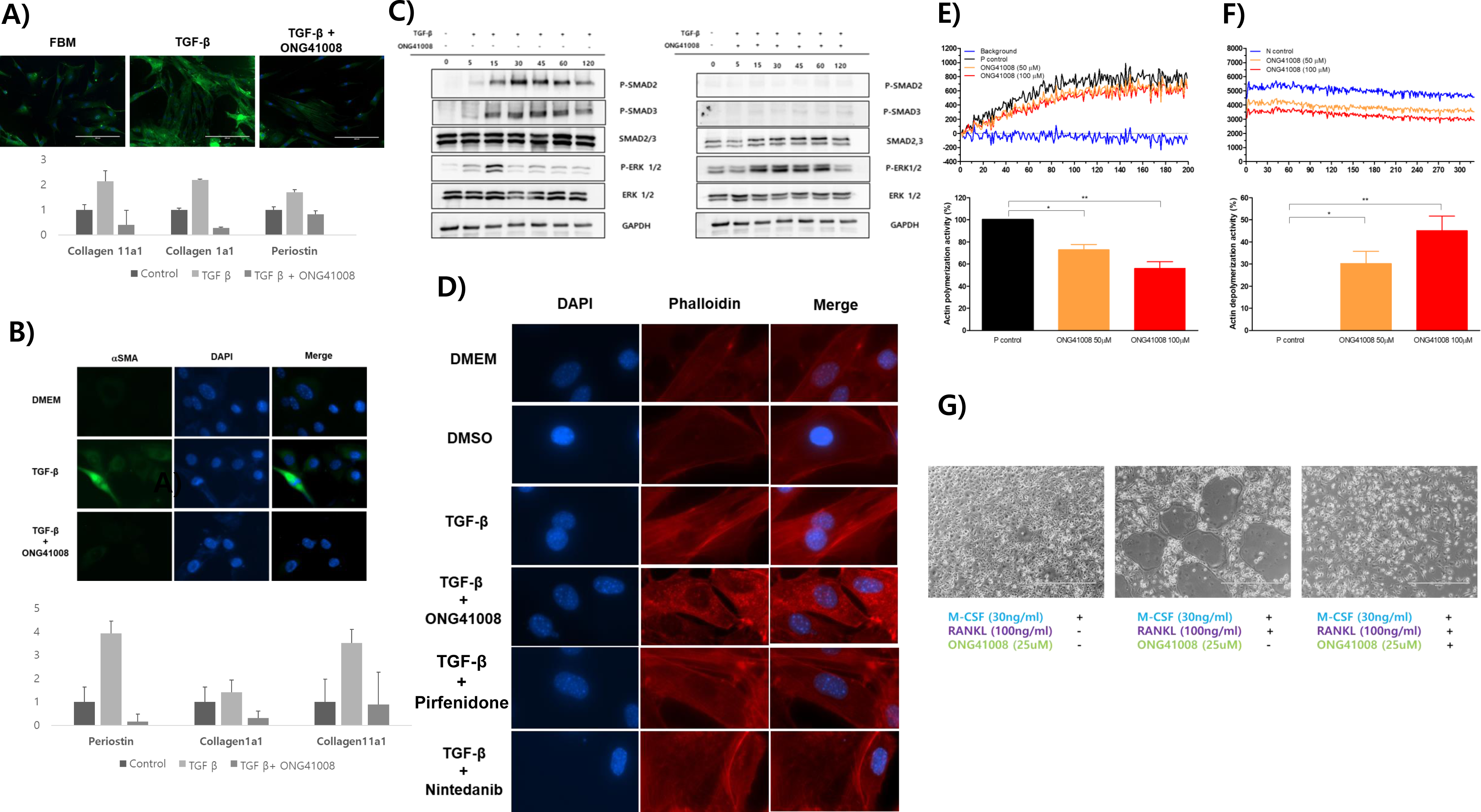
ONG41008 is a potent anti-fibrogenic inhibitor and a depolymerizer of F-actin. (A and B) ICC were performed for α-SMA on DHLF or ONGHEPA1 after 24hrs treatment with medium, TGFβ (2.5ng/ml), or TGFβ plus ONG41008 (25μM) and cell reactivities were observed under florescent microscope. *Periostin*, *Collagen1α1*, and *Collagen11α1* mRNA levels were measured by qPCR. (C) Kinetics of TGFβ-induced SMAD or ERK phosphorylation in presence of ONG41008 were conducted via western blot using the cell lysates of ONGHEPA1 treated with TGFβ in the presence or absence of ONG41008 at various time points; 0, 5, 15, 30, 45, 60, 120 mins. (D) Actin-phalloidin staining to test actin depolymerization was explored. ONGHEPA1 cells were treated with ONG41008, Pirfenidone, or Nintedanib in the presence of TGFβ as compared to control. (E-F) *In vitro* actin polymerization and depolymerization assays, respectively, were conducted. Actin polymerization activity and amount of depolymerization were measured in kinetic mode fluorometer at different concentration of ONG41008. Statistical significance was calculated by Student’s t-test. *, P < 0.05, **, P<0.01. (G) Mouse bone-marrow cells were subjected to differentiation into macrophages in the presence of M-CSF for 3 days. Macrophages were counted, replated, and stimulated with RAKNL for 5 days in the presence or absence of ONG41008. Multinucleated osteoclasts were observed with phase-contrast microscopy.

To further elucidate mode of action (MOA) for ONG41008, we stimulated DHLF or ONGHEPA1 with TGFβ plus ONG41008 and stained them with various antibodies recognizing the components of LTC (late TGFβ complex). LTC plays a central role of TGFβ biogenesis (15), releasing active TGFβ, which is then engaged in the nascent TGFRI/TGFRII complex, giving rise to phosphorylation of SMAD2/SMAD3. Among the known components, LTBP1 and LAP1 were tremendously induced and secreted into ECM (extra cellular matrix) compartments with the treatment of TGFβ, but LTBP4 appears to be either only partially affected or unaffected by ONG41008 in DHLF (Figure 2A). This data suggests that pathogenic myofibroblasts themselves may express LTC and may be becoming further pathogenic. ONGHEPA1 were also treated with TGFβ in the presence or absence of ONG41008 for 24 hr. It was evident that LTBP1, LAP1 and Integrin α5β3 in the presence of ONG41008 were massively downregulated and appeared to be localized in endosome-like compartments and actin stress fibers were substantially disintegrated (Figure 2B). Taken together, these combined data strongly indicate that ONG41008 is a noble TGFβ biogenesis inhibitor, thereby dismantling LTC and leading to cessation of TGFR signaling.

**Figure 2.**
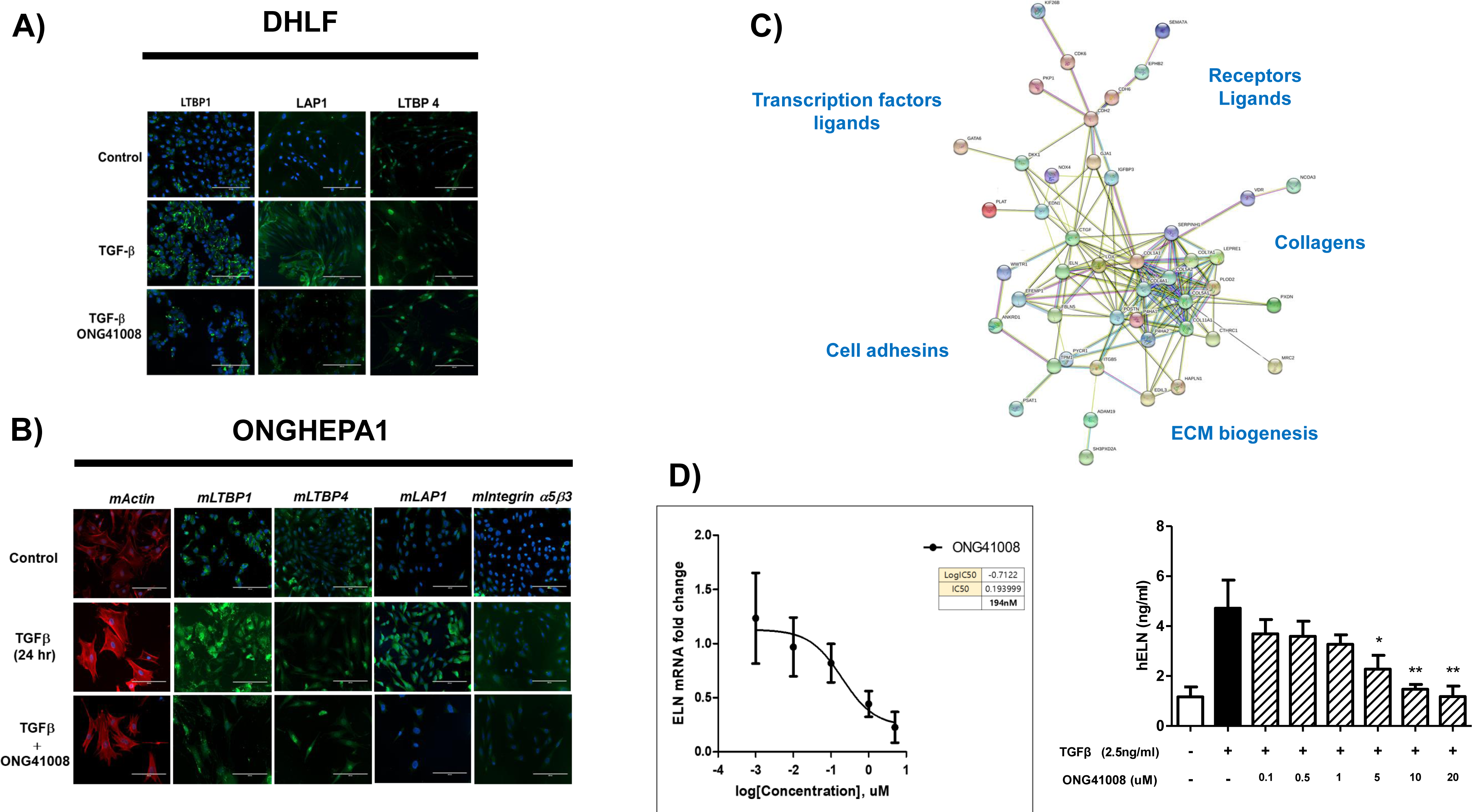
ONG41008 is a TGFβ biogenesis inhibitor via dismantling LTC. (A) DHLF were stimulated with TGFβ in the presence or absence of ONG41008 and stained for F-actin (phalloidin, red), LTBP1 (green), LAP1 (green), LTBP4 (green), and DAPI (blue). (B) ONGHEPA1 cells were stained for F-actin (phalloidin, red), mLTBP1 (green), mLTBP4 (green), mLAP1 (green), integrin α5β3 and DAPI (blue). C) Transcriptomic analysis revealed a set of genes, seventy-seven, called TGFβ-induced fibrosis-inducing genes (FIGS), which are significantly downregulated by ONG41008. An interactome study was conducted using the STRING database based on the above listed seventy-seven FIGS. D) Identification of ELASTIN as a potential pharmacodynamic marker associated with ONG41008-mediated anti- fibrosis. IC50 of ONG41008 for *Eln* mRNA expression was determined by qPCR. Culture supernatants from DHLF treated with TGFβ or TGFβ plus ONG41008 were examined for ELN protein levels by ELISA. **P < 0.001 and *P <0.05 relative to vehicle control were determined by Student’s t-tests.

### Generation of ONG41008-mediated fibrogenic interactome in DHLF and identification of potential pharmacodynamic markers

In order to identify the ONG41008-mediated fibrosis-inducing genes (FIGS), DHLF were differentially stimulated with TGFβ in the presence or absence of ONG41008 and transcriptomic change was analyzed by RNA-seq. Schematic representation of this selection process is shown in supplementary Figure 1 and a resulting interactome representing a fibrogenic proteome in DHLF is shown in Figure 2C. We nailed down seventy-seven FIGS whose p-values were below p>0.005. Many of these have been already known to be fibrogenic genes such as *CTGF* (connective tissue growth factor), *PERIOSTIN*, *LOX* (lysyl-oxidase), or *N-cadherin (CDH2)* (16–18). Elastin *(ELN)* gene expression has been appreciated in that ELN is profibrotic and represents progress of fibrosis. It is also known to be an EMT gene (19, 20). Clear indication is that the *ELN* gene expression was substantially induced with TGFβ and was knocked down with ONG41008 whose IC50 was 194 nM (Figure 2D). In line with this, secreted ELN from DHLF was significantly inhibited with ONG41008 (Figure 2D). ELN could be use a pharmacodynamic marker for ONG41008. In addition, real-time PCR revealed six more potential candidates; *PERIOSTIN, COL1α1, COL5α1, ACTA2, COL11α1, and COL6α3* (supplementary Figure 2). In particular, the expression changes of *COL5α1* were notable that could be a good candidate for a PD marker in response to ONG41008. In addition to the DHLF interactome, an ONGHEA1-ONG41008 interactome was established and a fibrogenic interactome was well represented in that an array of pioneering fibrogenic protein clusters involving collagens, ECM biogenesis, adhesins, differentiation, and enzymes denoted by each respective colored circle were identified. The TGFβ-ONG41008 interactome comprised sixty-one FIGS. Compared to the DHLF interactome, cytokines or enzymes appeared to play important roles in initiating and/or perpetuating fibrosis in the liver. This data suggests that ONG41008 would be pertinent to exploration of a wide range of tissue fibrosis.

NOX4, a NADPH oxidase, caught our attention because induction of NOX4 has been reported for production of pronounced reactive oxygen species (ROS) as well as for elicitation of several important signaling events like inflammasome activation and has been implicated in an important causative gene for fibrogenesis in IPF (21) (22). To scrutinize whether ONG41008 was able to suppress induction of NOX4, thereby downregulating ROS production, ONGHEPA1, which had been developed as an *in vitro* fibrogenesis model, were stimulated with TGFβ and induction kinetics of *NOX4* was analyzed at transcription levels along with *COL11α1* and *PERIOSTIN*. As shown in Figure 3A, *NOX4* mRNA was gradually induced by TGFβ and ONG41008 was effectively able to block the induction. Interestingly, ONG41008 was also able to block induction of *CCL2* and *CCL7* encoding macrophage chemoattractant proteins, suggesting that ONG41008 could modulate migration of macrophages and attenuate inflammation in the lung. Suppression of *NOX4* may be related to downregulation of innate immunity mounted by macrophages. Production of ROS was also significantly attenuated in ONGHEPA1, which was explored by ICC (Figure 3B). The same held true for DHLF (Figure 3C). Furthermore, ROS production was markedly reduced via dye staining (Figure 3D).

**Figure 3.**
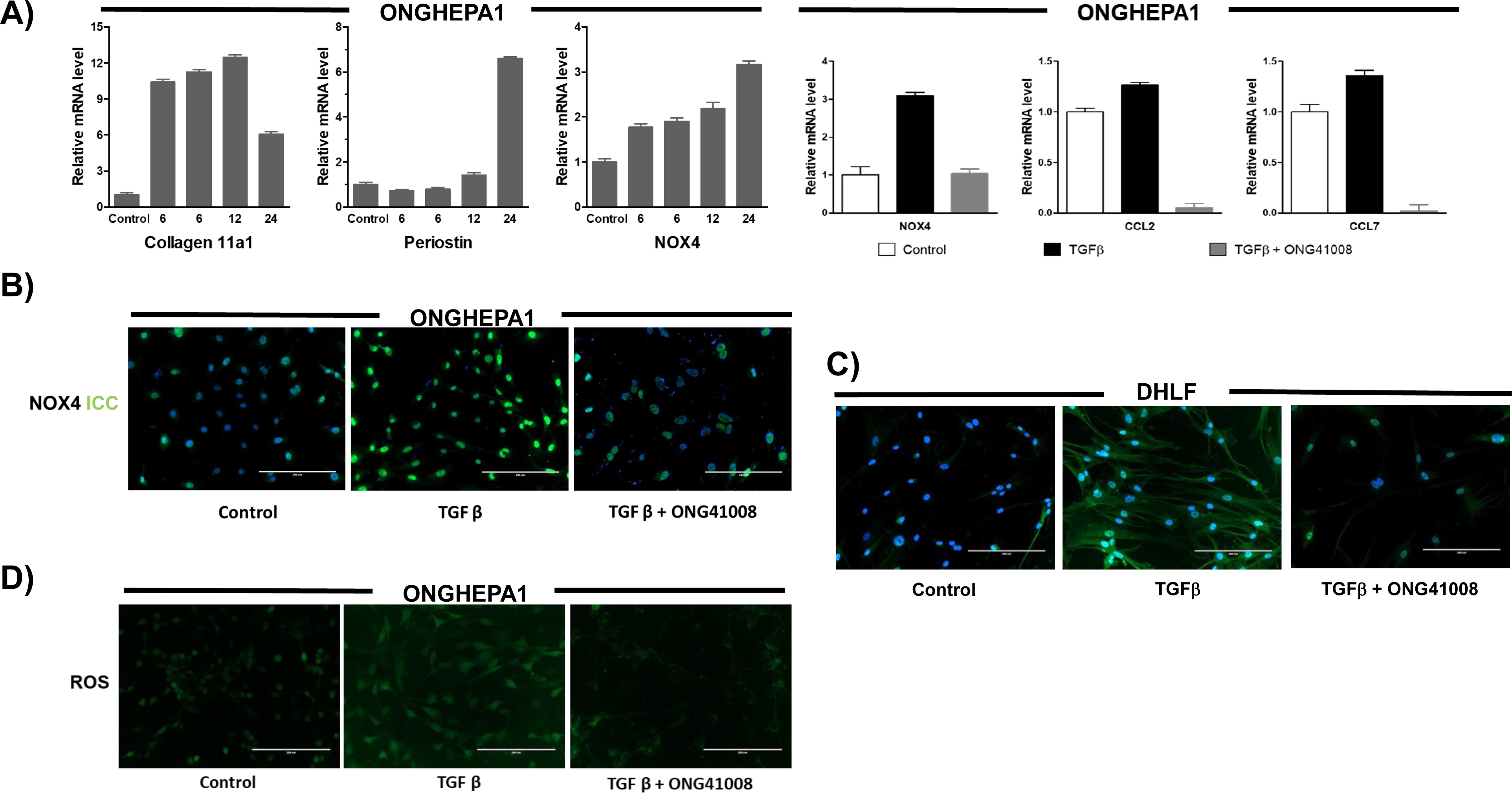
NOX4 is an anti-fibrogenic target by ONG41008. (A) qPCR for analysis of gene expression of Collagen*11α1, Periostin* and *NOX4* in TGFβ treated ONGHEPA1 was conducted at various time points; 0, 6, 9, 12, 24 hr. A set of qPCRs for *NOX4, CCL2, CCL7* in ONGHEPA1 was also performed. (B and C) ICC were performed for detecting NOX4 in ONGHEPA1 and DHLF, respectively, which had been incubated for 24hrs in control, TGFβ, or TGFβ plus ONG41008. (D) ROS assays were conducted with the use of DCFDA in ONGHEPA1.

Taken together, ONG41008 may exhibit a potent anti-fibrogenic capability in the lung by dismantling LTC and inhibitory effect on macrophage infiltration into fibrotic lesions may occur via modulation of NOX4 expression.

### ONG41008 was a potent inhibitor of innate immunity in that it was capable of blocking production of TNFα or chemokines affecting macrophages or T-cells in LPS-stimulated macrophages

It has been recently reported that knocking out *TNFα* leads to the lung or the liver inflammation exemplified by IPF, chronic obstructive pulmonary disease (COPD) or liver steatosis (23) (24). Only since we explored a potent anti-inflammatory capability associated with ONG41008 in HSC or DHLF in TGFβ-dependent manner, a natural question would be whether its anti-inflammatory property can be extended to macrophages in response to LPS stimulation in the absence of TGFβ. RAW264.7 cells, a mouse monocytic leukemic cell line resembling macrophages, were stimulated with LPS in the presence or absence of ONG41008. As shown in Figure 4A, expression of proinflammatory cytokines or chemokines including *TNFα*, *CCL2*, *CCL7*, *CXCL2* and *CXCL10* were markedly attenuated by ONG41008. Interestingly, *CHOP* or *Nox1* was so markedly reduced that formation of inflammasome or ancillary inflammatory signal transduction pathways could be affected (25) (26) (27). However, no statistically significant repression of *IL6* or *IL23* was made (data not shown). TNFα levels in the culture supernatants were clearly reduced with the treatment of ONG41008 (Figure 4B). An interactome representing the LPS-ONG41008 axis in macrophages was established via RNA-Seq. To our surprise, TNFα turned out to be the central nod of the interactome, strongly suggesting that ONG41008 may play a central role in antagonizing TNFα transcription (Figure 4B). The protein clusters involving chemokines, metabolism, innate immunity, and differentiation, whose important genes are denoted by blue circles, were downregulated by ONG41008 (Figure 4C) Summing up all these observations, ONG41008 was intrinsically equipped with both anti-fibrotic and anti-inflammatory capabilities, especially suggesting that the latter could suit regulation of innate immunity.

**Figure 4.**
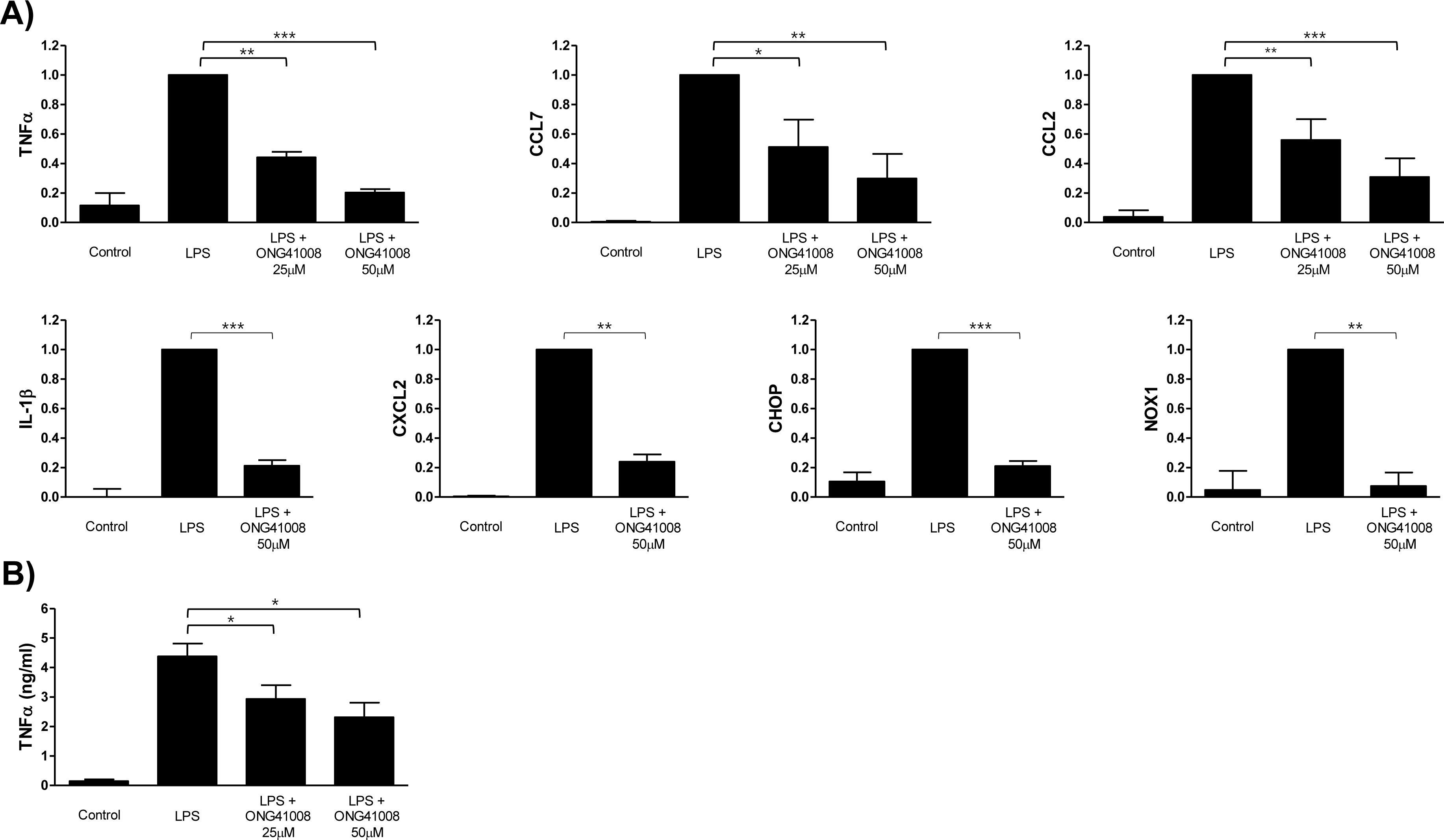
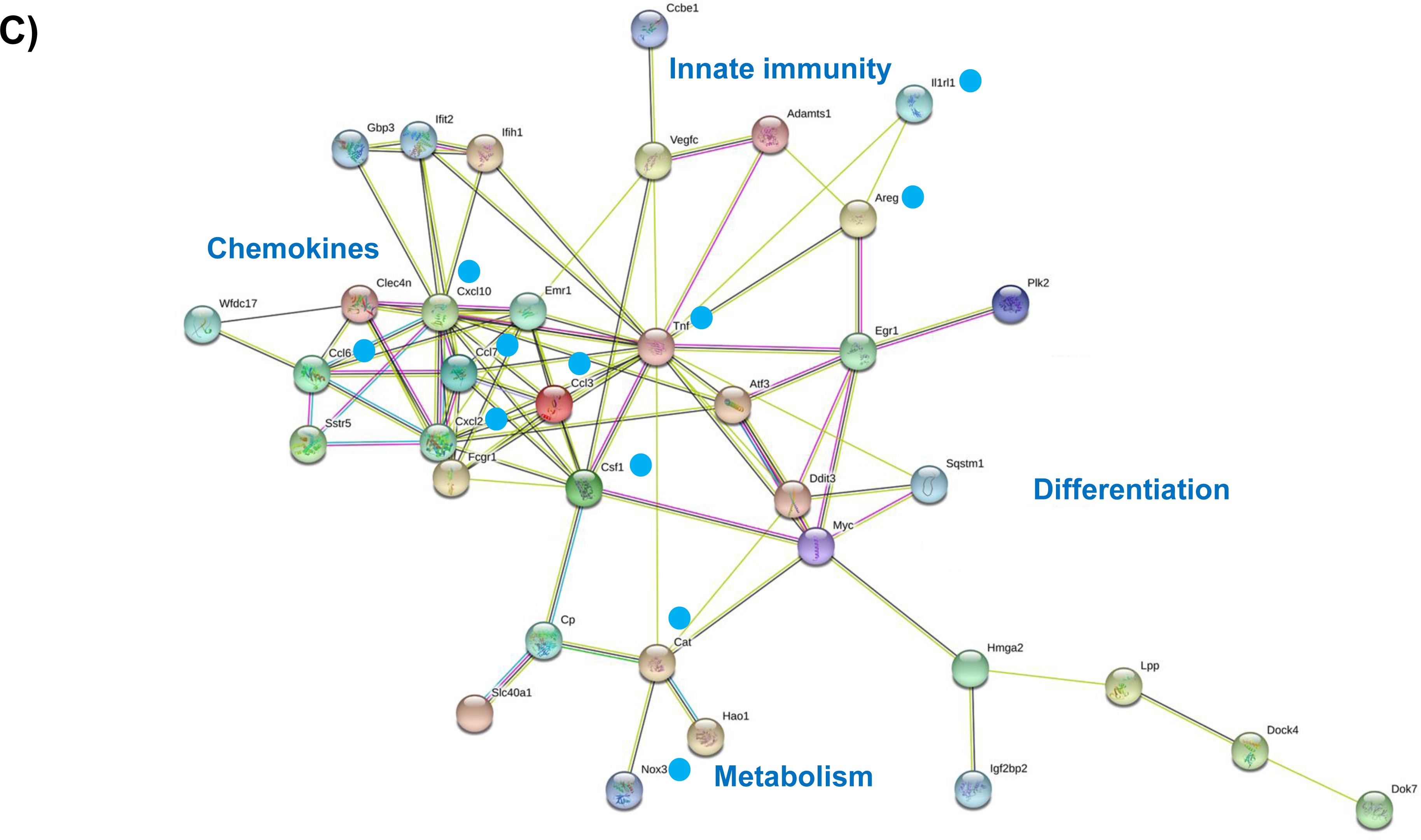
Inhibition of proinflammatory or inflammatory cytokines by ONG41008 in LPS-stimulated RAW264.7 cell. A) RAW264.7 cells were stimulated with LPS (100ng/ml) or LPS plus ONG41008 (25μM, 50μM) for 24hrs and mRNA expression for inflammatory markers were measured by qPCR. B) Supernatants of cultured RAW264.7 treated with LPS and/or ONG41008 were collected and ELISA was performed to measure the expression of TNFα. Data are presented as the mean ± standard deviation (n=3).

### *In vivo* therapeutic efficacy of ONG41008 for lung fibrosis

Prophylactic and therapeutic efficacy of ONG41008 for lung fibrosis were analyzed via BLM. Lower concentrations of ONG41008 or pirfenidone formulated in 4% HPβCD (2- hydroxypropyl)-beta-cyclodextrin) were used for BLM such that ONG41008 25mpk and 50mpk were able to significantly attenuate production of soluble collagen with statistics; 42% and 62% inhibition, respectively (Figure 5A). No decent inhibitory capacities associated with pirfenidone 100mpk or ONG41008 10mpk was noted. Only while significant inhibition of HP production was noted at ONG41008 50mpk with strong statistical significance reduction tendency was observed at both ONG41008 10mpk and 25mpk and at pirfenidone 100mpk (Figure 5B). We observed that pirfenidone consistently showed tendency in lowering TGFβ production (Figure 5C). To our surprise, the short-lasting PK tests prior to sacrifice of treated mice indicated that at least, 10-fold difference in plasma or lung tissue PK values existed between pirfenidone 100mpk and ONG41008 50mpk, strongly suggesting although bioavailability of these drugs exhibited a massive difference; pirfenidone 70∼80% vs ONG41008 1∼1.5%, nevertheless, the disease-correction efficacy of ONG41008 may be exceedingly higher than that of pirfenidone (Figure 5D). A separate BLM was conducted by a 3^rd^ party investigator in order to compare tissue staining for collagen and αSma. ONG41008 at those three administration concentrations remarkably reduced collagen deposition shown by Masson-Trichome tissue staining. Dexamethasone was able to block collagen deposition (Figure 5E). Expression of αSMA was more significantly decreased in all ONG41008-treated mice than control mice. However, dexamethasone did not affect αSMA staining, indicating that anti-inflammation may not be able to mitigate proliferation of myofibroblasts at advanced stages (Figure 5F).

**Figure 5.**
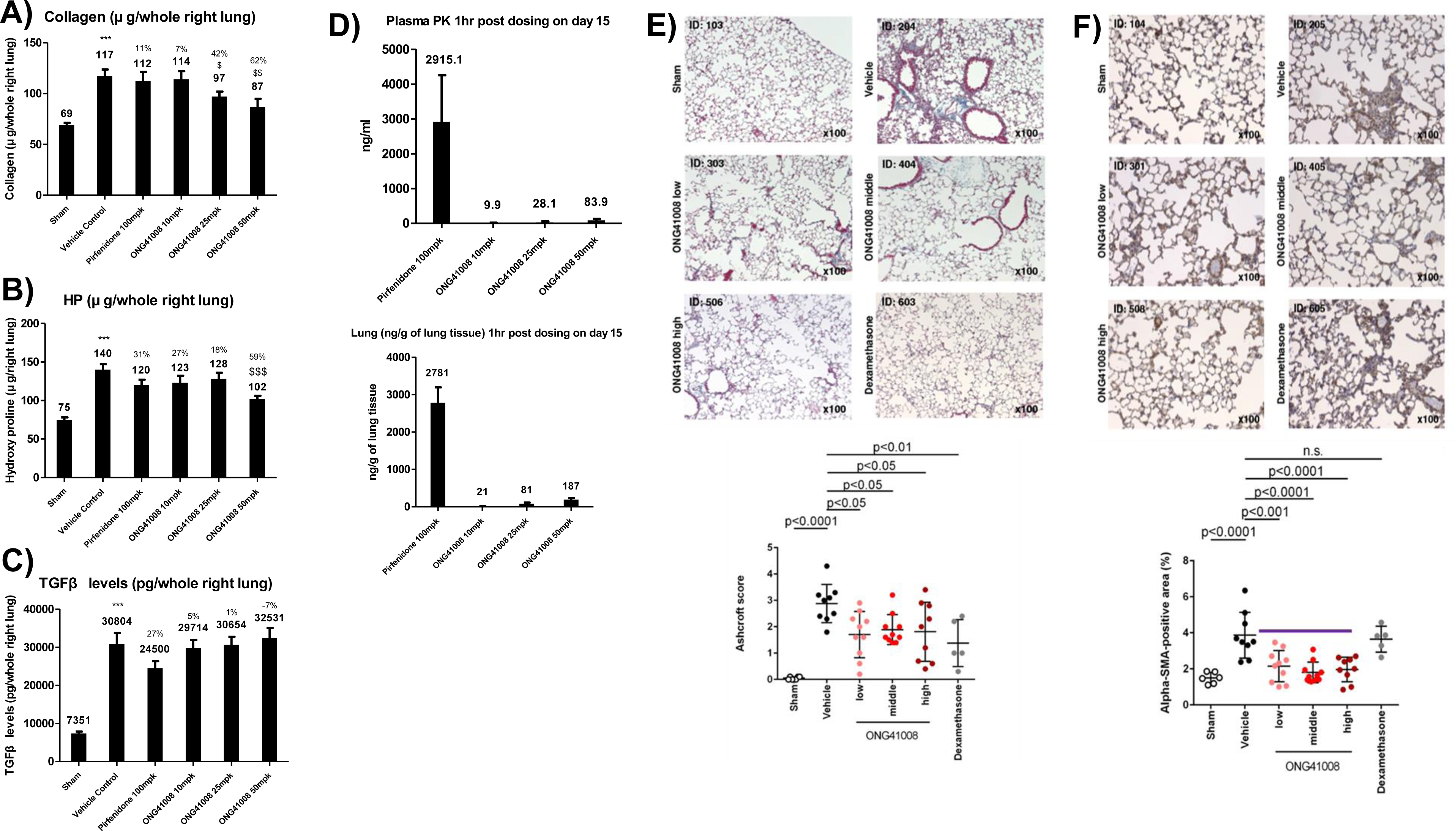
ONG41008 had positive effects against lung fibrosis in BLM prevention model. (A, B and C) Lung collagen, hydroxyproline and TGFβ levels measured by ELISA after treatment with pirfenidone or 10, 25, or 50mpk ONG41008 in a BLM prevention model. D) Plasma samples were collected 1hr post dosing on day 21 from BLM prevention model treated with Pirfenidone or 10, 25, or 50mpk ONG41008 were analyzed using LC-MS/MS analytical method. For lung samples analysis, right lung tissue homogenates collected 1hr post dosing on day 21 was analyzed using an LC-MS/MS analytical method. Statistical significance was calculated by Student’s t-test. ***, P < 0.001, relative to Sham. $, P < 0.05, $$$, P < 0.001 relative to vehicle control.

Therapeutic BLM was explored. As shown in Figure 6A, no significant inhibition of collagen production was noted at ONG41008 10mpk and 25mpk as well as at pirfenidone 100mpk but decreasing tendency between 26% and 36% remained certain, suggesting that larger exposure of ONG41008 may be needed to attenuate existing the lung fibrosis in the therapeutic model. Indeed, ONG41008 50mpk significantly inhibited collagen production with a statistical significance. Significant inhibitions of HP production were accomplished at both ONG41008 25mpk and 50mpk, 37% and 34%, respectively. Although pirfenidone 100mpk exhibited some tendency of reduction in HP production, however, it did not reach statistical significance (Figure 6B). No ONG41008 10mpk did affect HP production. Interestingly, a set of histopathology analysis using collagen morphometry revealed all regimens representing pirfenidone 100mpk and the varying concentrations of ONG41008 gave rise to comparable and statistically significant inhibition of collagen deposition with average 40%, indicating that both drugs may be able to ameliorate the established lung fibrosis (Figure 6C). While pirfenidone 100mpk resulted in average 31% reduction in TGFβ production in the lung lysates with or without statistical significances (data not shown) no ONG41008 influenced TGFβ production (Figure 6D). Consistent with the prophylactic data on the final PK tests before sacrifice the bioavailability of pirfenidone 100mpk in both plasma and lung lysates were at least ten times higher than that of ONG41008 50mpk, however, disease-correction efficacy of ONG41008 25mpk or 50mpk was minimally equivalent to that of pirfenidone 100mpk but at best was better that of pirfenidone 100mpk (Figure 6E and 6F).

**Figure 6.**
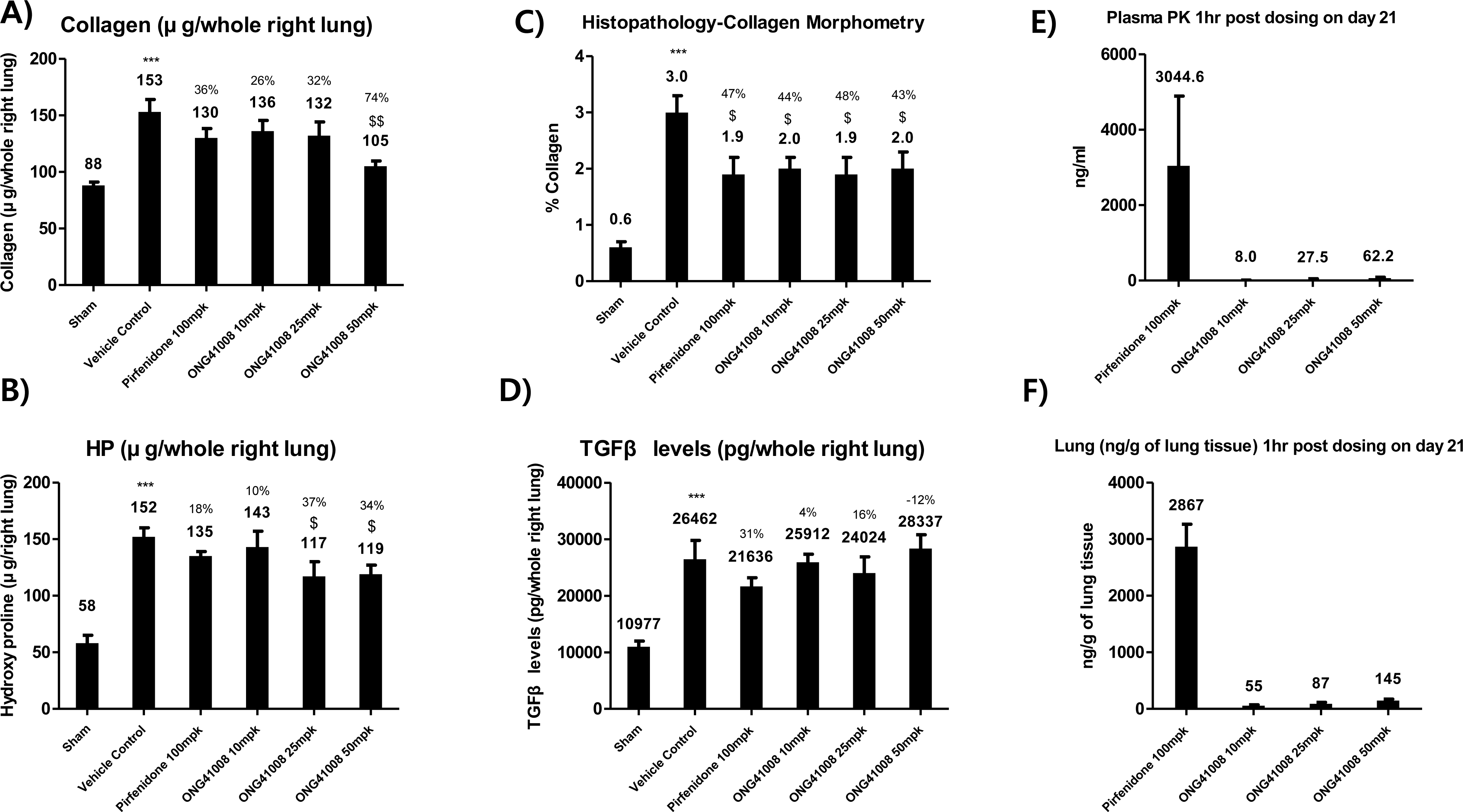
Anti-fibrotic effect of ONG41008 in a therapeutic bleomycin-induced lung fibrosis model. (A, B and D) Lung collagen, hydroxyproline and TGFβ levels were measured by ELISA after pirfenidone or different dosages of ONG41008 (10, 25, 50mpk) oral administration in BLM therapeutic model. Percent inhibition relative to Vehicle control are presented with average measurement. (C) Lung sections were analyzed by Picro-Sirius red staining for morphometric analysis. Random field photographs were taken under 20x magnifications for % collagen and expressed as percentage area. E) Plasma samples collected 1hr post dosing on day 21 from BLM therapeutic model treated with Pirfenidone or 10, 25, or 50mpk ONG41008 were analyzed using LC-MS/MS analytical method. F) Right lung tissue homogenates collected 1hr post dosing on day 21 was analyzed using an LC-MS/MS analytical method. Statistical significance was calculated by Student’s t-test. ***, P < 0.001, relative to Sham. $, P < 0.05, $$$, P < 0.001 relative to vehicle control.

Taken together, ONG41008 could be a good candidate for treating IPF as a monotherapy or as a combination therapy with pirfenidone.

## Discussion

CSD have been well appreciated for their anti-inflammatory capabilities (28). We recently found that a few CSD are capable of attenuating *in vitro* fibrogenesis and *in vivo* fibrosis (29). Slight modification of functional groups linked to CSD severely affected degree of anti-fibrogenesis, indicating that medicinal chemistry would be able to improve their therapeutic efficacy. One significant drawback associated with CSD in terms of *in vivo* use for treating fibrosis is that CSD largely tend to undergo glucuronidation. We confirmed that glucuronide form of CSD completely lost anti-fibrogenic capability *in vitro* (data not shown). Thus, to cope with this limitation, development of new CSD analogs should have been implemented. ONG41008 rendered non-toxic when subjected to 14 Day’s oral gavage non-GLP toxicity tests. PK profiles using mice and rats or beagle dogs via oral administration were readily detectable in plasma and lung. Therefore, we concluded that ONG41008 could be a therapeutically pertinent drug candidate.

The major MOA seems to dismantle LTC in such a way that ONG41008 limits binding of active TGFβ to TGFRs; one major manifestation is blocking phosphorylation of SMAD2/SMAD3, resulting in disabling type 2 EMT called fibrogenesis. However, phosphorylation of ERK was not affected by ONG41008, suggesting that ONG41008 is largely associated with the main signaling of TGFR, i.e. phosphorylation of SMAD2/SMAD3. Crucial role of TNFα associated with lung and liver fibrosis prompted us to scrutinize effect of ONG41008 on macrophage physiology. Macrophages play a central role in inflammation whose activation has been known to be in part modulated by ROS (30) (31). As ONG41008 mitigated *NOX4* gene expression in DHLF as well as in ONGHEPA1, how ONG41008 regulates *NOX4* gene expression in these cells remains intriguing. ONG41008 was able to completely inhibit transcription of *NOX4* as well as NOX4 expression in conjunction with *COL11α1* or *PERIOSTIN*. According to an interactome analysis shown in Figure 2C, NOX4 seems to be functionally related to endothelin (EDN) 1, connective tissue growth factor (CTGF) and insulin-like growth factor binding proteins (IGFBP) 3. This data suggests that NOX4 plays an important role in pathogenesis of myofibroblasts. ONG41008 seemed able to block innate immunity in macrophages via antagonizing transcription of *TNFα* . These two-combined acting mechanisms, anti-fibrotic and anti-inflammatory capabilities, may be responsible for efficiently disarming transdifferentiation of progenitors into myofibroblasts or proliferation of myofibroblasts, resulting in mitigation of collagen or hydroxyproline production. Anti-inflammatory capability associated with ONG41008 may play an important role in giving rise to beneficial features for amelioration of IPF. It could initially prevent fibrogenic presumably provided by the lung stromal cells and promote cellular homeostasis such as cell survival. Based on these observations, we propose a working hypothesis on MOA associated with ONG41008 in the first line of defense (Figure 7).

**Figure. 7.**
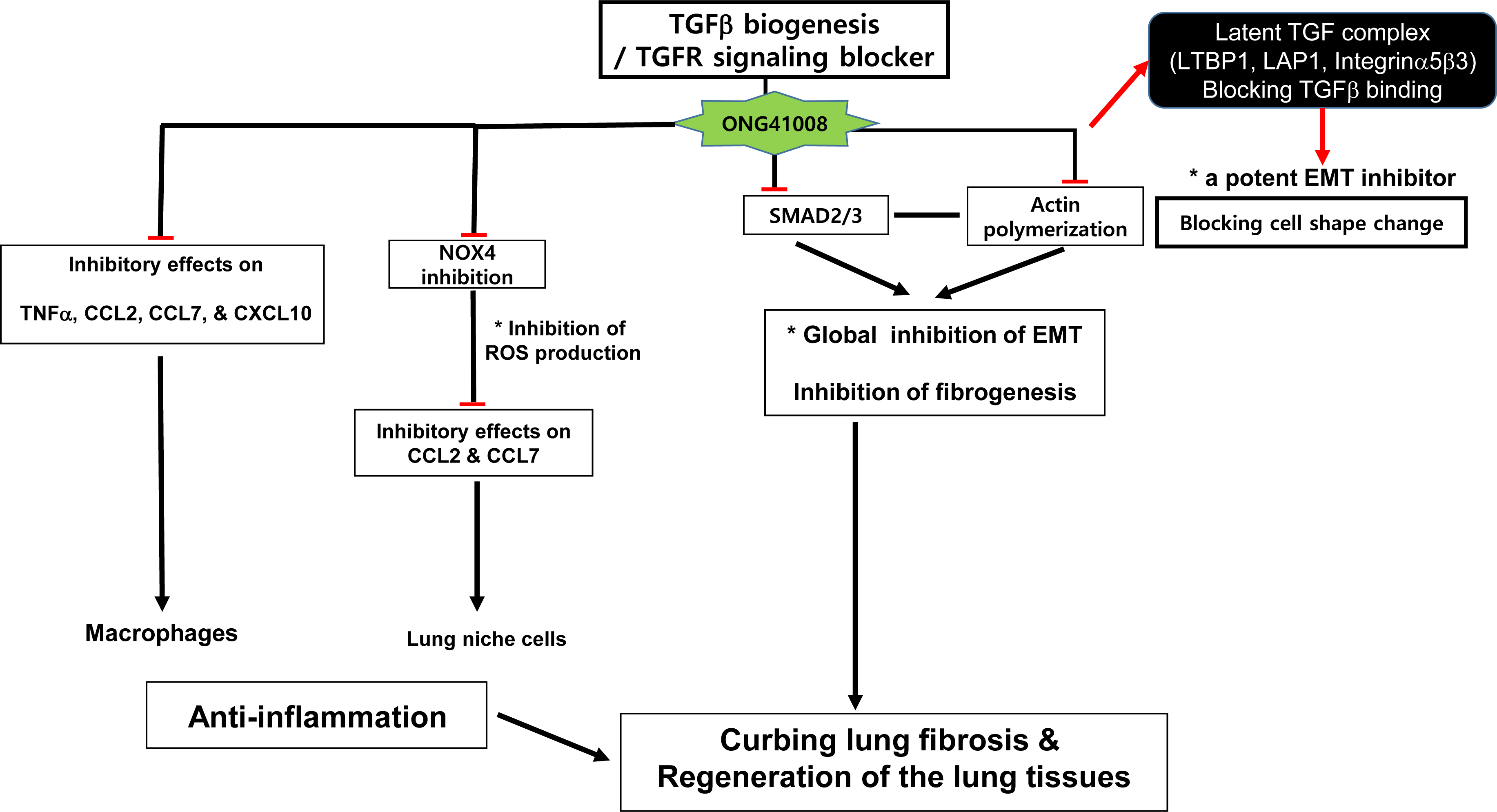
Potential mechanism of ONG41008 action in the treatment of organ fibrosis. Major acting anti-fibrotic mechanism associated with ONG41008 disintegrates actin stress filaments via depolymerization of F-actin, leaving latent TGFβ complex dismantled. Inability of TGFβ binding to TGFβ II or I signaling receptor is unable to generate canonical or noncanonical TGFR signaling leading to attenuation of EMT. Furthermore, ONG41008 blocks phosphorylation of SMAD2/3. These two acting mechanisms may be responsible for efficiently disarming proliferation of or collagen or hydroxyproline production of myofibroblasts. Inhibitory effects on *NOX4* expression or macrophage activation may be responsible for anti- inflammation, leading to homeostasis.

Taken together, we generated a CSD called ONG41008 having both anti-fibrotic and anti-inflammatory capabilities. ONG41008 is able to ameliorate a broad range of fibrotic diseases such as cardiovascular fibrosis, kidney fibrosis, NASH or fibrotic cancers such as pancreatic cancer.

## Acknowledgments

We greatly appreciate Dr. HS Yoon (NTU Singapore) for scientific advices. A special thank is given to Dr. JB Kim for useful experimental inputs. We are thankful to D Lee (being the CTO in Osteoneurogen) for proof reading the manuscript and to all Osteoneurogen researchers and administrative workforces who have been helping us, making the current manuscript possible. This study was supported by an intramural fund from Osteoneurogen, Inc. (ONG-400022)

## Author contributions

B-S Youn and H-S Kim conceived the idea and composed the manuscript. I-H Kim and B L. Seong exchanges the ideas related to therapeutic efficacy of ONG41008 to fibrotic diseases. MK Maeng was largely involved in executing major experiments as well as organizing and sorting out all involved methods and materials. JY Lee played a role in TGFβ signaling along with ONG41008. SB Kim played a major role in trans-differentiation of HSC into pathogenic myofibroblasts. The major body of this manuscript has been initially exposed to public on the bioRxiv depository since September, 2019 (doi: https://doi.org/10.1101/770404).

## Animal care

Mice, rats or beagle dogs were managed by the IACUC associated with Syngene International (Bangalore, India).

## Footnotes

The following abbreviated terms were mainly used in this manuscript;

**CS**: Chromone Scaffold

**CSD**: Chromone Scaffold Derivatives

**IPF**: Idiopathy Pulmonary Fibrosis

**DHLF**: Diseased Human Lung Fibroblasts from IPF patients

**HSC**: Hepatic Stellate Cells

**BLM**: Bleomycin-Induced Lung Fibrosis **NASH**:Non-Alcoholic Steato Hepatitis **LTC**: Latent TGF Complex

**HP**: Hydroxyprolie

**MOA**: Mode of Action

**FIGS**:fibrosis-inducing genes

## Materials and Methods

### Cell culture and reagents

DHLFs were purchased from Lonza (Basel, Switzerland) and cultured in fibroblast growth medium (FBM, Lonza, Walkersville, MD, USA). Recombinant human TGFβ and PDGF were obtained from Peprotech (Rocky Hill, CT, USA) and used at a final concentration of 5 ng/ml. Chemically synthesized ONG41008 was obtained from Syngene International Ltd. (Bangalore, India), dissolved at a stock concentration of 50 mM in DMSO, and stored in aliquots at -20°C. DMSO with according concentration was used as control. RAW264.7 cell line was purchased from Korean Cell Line Bank (Seoul, Korea) and cultured in RPMI supplemented with 10% FBS and 1% P/S (Welgene, Seoul, Korea). LPS was purchased from Sigma and used at final concentration of 100 ng/ml.

### Induction of osteoclast differentiation

Animal experimental protocols were approved by the institutional Animal Care and Use Committee at Yonsei University Wonju College of Medicine (Identification code: YWC- 200325-1) and procedures were performed in accordance with the guidelines of the National Institutes of Health’s Guide for the Care and Use of Laboratory Animals. Bone marrow mononuclear cells and marrow-derived macrophages were derived as previously described (35). Briefly, mouse bone marrow cells were harvested from femurs and tibias from 8-week-old Balb/c mice and cultured overnight on 100-mm dishes in α-MEM )(WelGENE Inc., Republic of Korea) supplemented with 10% fetal bovine serum (FBS) and penicillin/streptomycin. Floating cells were collected and further cultured in the presence of M-CSF (30 ng/ml) for 4 days to generate bone marrow-derived macrophages (BMM). To induce osteoclast differentiation, BMM were stimulated with 100 ng/mL RANKL and 30 ng/mL M-CSF in the presence or absence of ONG41008.

### Actin polymerization/depolymerization assay

Actin Polymerization/Depolymerization Assay Kit (Abcam, ab239724) was used for both assays. For actin polymerization assay mixture of; actin, buffer, and test samples were placed in 96 plate well in order of supplemented Buffer G (2μM ATP, 5μM DTT), actin, and test samples. As test samples, Buffer G was used as background control, DMSO as positive control, and ONG41008 in DMSO with concentration range from 25 to 100μM were used. The mixtures in each well were then pipetted thoroughly to mix and incubated in dark, at room temperature, for 15 min. After incubation the polymerization activation buffer, supplemented 10X Buffer P (10mM ATP) was added into each well and thoroughly pipetted. For actin depolymerization assay; Buffer G (2μM ATP, 5μM DTT), supplemented 10X Buffer P (10mM ATP), and actin were placed in 96 plate well in listed order and pipetted to mix thoroughly. The mixture was incubated in dark, at room temperature, for 1 hr for polymerization to take effect. After the incubation test samples, DMSO as negative control and ONG41008 at concentration from 25 to 100μM, were added to each appropriate well and pipetted thoroughly. Florescence of both actin polymerization and depolymerization assay were measured with Florescence/Luminescence Analyzer Hidex Sense at florescence Ex/Em 355/405nm (range 5∼10nm) in kinetic mode for 2 hr (180 cycles).

### Reverse transcriptase PCR and real-time PCR

Cells cultured in either 12 or 24-well plates were washed twice with cold PBS and harvested using TaKaRa MiniBEST Universal RNA extraction kit (Takara, Japan). RNA was purified using the same kit according to manufacturer’s protocol. RNA was reverse-transcribed using the cDNA Synthesis Kit (PCRBio Systems, London, UK). Synthesized cDNA was amplified with StepOne Plus (Applied Biosystems, Life Technologies) and 2× qPCRBio Probe Mix Hi-ROX (PCRBio). Comparisons between mRNA levels were performed using the ΔΔCt method, with GAPDH as the internal control.

### ELISA and immunoblotting

Mouse TNFα Quantikine ELISA kit was purchased from R&D systems (Minneapolis, MN, USA). Raw264.7 cell was seeded at 1 x 10^5^ in 12 well cell culture plate and incubated O/N. The cells were then treated with LPS (100ng/ml) and ONG41008 at the indicated concentrations. Balancing amount of DMSO was added to each treatment condition. After 24HRs of treatment supernatant from each was collected for ELISA. The assay was performed according to the manufacturer’s manual. For three repeat experiment standard curve was made each time to calculate TNFα concentration. DHLF at a density of 2x10^5cells/ml were seeded onto 24well plate, followed by treatment with TGFβ (2.5ng/ml) and TGFβ (2.5ng/ml) plus various concentrations of ONG41008. After 24 hr, the supernatants were collected and performed with human elastin ELISA kit (Abcam, ab239433) following the manufacturer’s protocol. For western blotting, antibodies for CD14, MD-2, P62, GAPDH were purchased from Abcam (Cambridge, UK), MyD88, TLR4 from R&D systems, NOX4 (14347-1-AP) from Proteintech (Rosemont, IL, USA), and LC3 A/B, pSMAD-2, pSMAD-3, SMAD2/3, pERK 1/2, ERK 1/2 from Cell Signaling Technology (Danver, MA, USA). All antibodies were diluted to 1:1000 v/v in 5% BSA in DPBS (Welgene).

### Immunocytochemistry

Cells were fixed using 4% paraformaldehyde, permeabilized with 0.3% TritonX100, blocked and incubated with 1:500 anti α-Smooth Muscle Actin (Young In Frontier, Korea), 1:500 Actin Phalloidin (Thermo Fisher, USA), 1:500 anti-LTBP1 (Aviva Systems Biology, San Diego, USA), 1:500 anti- LTBP4 (Aviva Systems Biology), 1:500 anti-LAP1 (Abcam), 1:500 anti-integrin α5β3 (biorbyt, Cambridge, UK), 1:500 anti-NOX4 (Proteintech), 1:500 anti-CD14 (Abcam), 1:500 anti-MyD88 (R&D systems), 1:500 anti-TLR4 (R&D systems) or 1:500 anti-MD-2 (Abcam) antibodies and 1:200 FITC conjugated secondary antibody (Young In Frontier) and imaged with fluorescence microscope EVOS® FL (Thermo). Nuclei were stained with DAPI

### Confocal imaging

Cells were fixed with 4% paraformaldehyde, permeabilized with 0.4% Triton X100, blocked with 1% BSA and incubated with anti-CD14 (1:500, Abcam), anti-Myd88 (1:500, R&D systems) for overnight at 4°C. After FITC conjugated secondary antibody incubation (1:200, Ab Frontier) for 1 hr at 37°C, images were acquired using Laser scanning confocal microscope (Carl Zeiss LSM700) with 63x oil immersion lens. Confocal images were analyzed with Zen black software.

### ROS staining

Intracellular production of ROS was detected after stimulation of ONGHEAPA1 cells with TGFβ (5ng/ml) or TGFβ plus ONG41008 (50μM). Cells were incubated with 10μM H2DCFDA (Abcam) for 1 hr in dark at 37°C. The cells were then examined with fluorescence microscope EVOS® FL (Thermo).

### RNA-seq processing, differential gene expression analysis, and interactome analysis

Processed reads were mapped to the *Mus musculus* reference genome (Ensembl 77) using Tophat and Cufflink with default parameters. Differential analysis was performed using Cuffdiff using default parameters. Further, FPKM values from Cuffdiff were normalized and quantitated using the R Package Tag Count Comparison (TCC) to determine statistical significance (e.g., P values) and differential expression (e.g., fold changes). Gene expression values were plotted in various ways (i.e., Scatter, MA, and Volcano plots), using fold-change values, using an R script developed in-house. The protein interaction transfer procedure was performed using the STRING database with the differentially expressed genes. A 60 Gb sequence was generated, and 10,020 transcripts were read and compared. The highest-confidence interaction score (0.9) was applied from the *Mus musculus* species, and information about interacts were obtained based on text mining, experiments, and databases (http://www.string-db.org/). Due to company information sake the above detailed RNA-Seq or interactome data interpretation would be limited but essential data sufficiently supporting our assertion were provided.

### Effects of drugs on bleomycin-induced lung tissue fibrosis

C57BL/6J mice were anesthetized by inhalation of 70% N2O and 30% O2 gas containing 1.5% isoflurane. Fifty microliters of bleomycin solution in distilled water was directly injected into the lungs, all at once, via the aperture. Immediately after injection, the mice were allowed to recover from the anesthetic, and then housed in normal cages. Bleomycin (0.03U BLM in 50µl saline) was administered once using a visual instillobot. Twelve days after the administration of bleomycin, ONG41008 was forcibly nasally administered via a micropipette, once a day (five times a week) for 1 week. ONG41008 was dissolved in water containing 4% 2-hydroxypropyl)-Beta-Cyclodextrin, and 1mpk was administered based on the most recent body weight. For 2 to 3 days after administration of ONG41008, mice were monitored for toxic symptoms or death, but no abnormal symptoms were observed. Three mice per test group were selected, and their lung tissues were excised. The lung tissues were stained with Masson’s trichrome and observed under a microscope. The degree of fibrosis of the lungs was assessed by an independent pathologist using the Ashcroft scoring system. Results were expressed as mean values and standard deviations. One hour before sacrifice, a final dose of ONG41008 or pirfenidone was administered for plasma or lung PK. The bleomycin-treated mice exhibited a rapid decline in weight, but the sham control behaved normally. ONG41008- or pirfenidone-administered mice exhibited weight gain from day 3 onward. Control and ONG41008-treated mice data were compared using Student’s *t*-test. “Differences between samples were considered statistically significant when p<0.05.

## Supplemental information

**Supplementary Figure 1. Schematic presentation of fibrosis-inducing gene selection process and TGFβ-ONG41008 interactome.** Transcriptomic analysis in DHLF treated with TGFβ or TGFβ plus ONG41008 was conducted. Differential expression was assessed with p< 0.005.

**Supplementary Figure 2. Selection of potential PD markers in DHLF stimulated with ONG41008 in the presence of TGFβ.** Real-time PCR experiments were performed with gene-specific primer sets and quantitative data were acquired. Gene expression changes were assessed with denoted p-values; 1) * corresponds to p<0.1 and ** corresponds to p<0.05.

**Supplementary Figure 3. Transcriptomic analysis and establishment of a ONG41008- specific interactome in ONGHEPA1** RNA-seq was performed by using total RNAs from ONGHEPA1 treated by control, TGFβ, or TGFβ + ONG41008. Differential expression was explored with p< 0.005. An interactome study using STRING based on sixty-one FIGS derived from RNA-Seq. Representative interactomes are denoted by colored ovals.

## References

1. Meyer KC. Pulmonary fibrosis, part I: epidemiology, pathogenesis, and diagnosis. Expert Rev Respir Med 2017; 11: 343–359.

2. Raghu G, Richeldi L, Jagerschmidt A, Martin V, Subramaniam A, Ozoux ML, Esperet CA, Soubrane C. Idiopathic Pulmonary Fibrosis: Prospective, Case-Controlled Study of Natural History and Circulating Biomarkers. Chest 2018; 154: 1359–1370.

3. Kramann R, Schneider RK. The identification of fibrosis-driving myofibroblast precursors reveals new therapeutic avenues in myelofibrosis. Blood 2018; 131: 2111–2119.

4. Ignat SR, Dinescu S, Hermenean A, Costache M. Cellular interplay as a consequence of inflammatory signals leading to liver fibrosis development. Cells 2020; 9: E461.

5. Lemoinne S, Friedman SL. New and emerging anti-fibrotic therapeutics entering or already in clinical trials in chronic liver diseases. Curr Opin Pharmacol 2019; 49: 60–70.

6. Miao CM, Jiang XW, He K, Li PZ, Liu ZJ, Cao D, Ou ZB, Gong JP, Liu CA, Cheng Y. Bone marrow stromal cells attenuate LPS-induced mouse acute liver injury via the prostaglandin E 2-dependent repression of the NLRP3 inflammasome in Kupffer cells. Immunol Lett 2016; 179: 102–113.

7. Dixon RA, Pasinetti GM. Flavonoids and isoflavonoids: from plant biology to agriculture and neuroscience. Plant Physiol 2010; 154: 453–457.

8. Stapleton AE, Walbot V. Flavonoids can protect maize DNA from the induction of ultraviolet radiation damage. Plant Physiol 1994; 105: 881–889.

9. Leyva-López N, Gutierrez-Grijalva EP, Ambriz-Perez DL, Heredia JB. Flavonoids as Cytokine Modulators: A Possible Therapy for Inflammation-Related Diseases. Int J Mol Sci 2016; 17.

10. Emami S, Ghanbarimasir Z. Recent advances of chroman-4-one derivatives: synthetic approaches and bioactivities. Eur J Med Chem 2015; 93: 539–563.

11. Gacche RN, Meshram RJ, Shegokar HD, Gond DS, Kamble SS, Dhabadge VN, Utage BG, Patil KK, More RA. Flavonoids as a scaffold for development of novel anti-angiogenic agents: An experimental and computational enquiry. Arch Biochem Biophys 2015; 577–578: 35-48.

12. Kim JY, Lee MS, Baek JM, Park J, Youn BS, Oh J. Massive elimination of multinucleated osteoclasts by eupatilin is due to dual inhibition of transcription and cytoskeletal rearrangement. Bone Rep 2015; 3: 83–94.

13. Troeger JS, Mederacke I, Gwak GY, Dapito DH, Mu X, Hsu CC, Pradere JP, Friedman RA, Schwabe RF. Deactivation of hepatic stellate cells during liver fibrosis resolution in mice. Gastroenterology 2012; 143: 1073–1083.e1022.

14. Wang Y, Brooks PJ, Jang JJ, Silver A, Arora PD, McCulloch CA, Glogauer M. Role of actin filaments in fusopod formation and osteoclastogenesis. Biochim Biophys Acta 2015; 1853: 1715–1724.

15. Robertson IB, Horiguchi M, Zilberberg L, Dabovic B, Hadjiolova K, Rifkin DB. Latent TGF-β-binding proteins. Matrix Biol 2015; 47: 44–53.

16. Makarev E, Izumchenko E, Aihara F, Wysocki P, Zhu Q, Buzdin A, Sidransky D, Zhavoronkov A, Atala A. Common pathway signature in lung and liver fibrosis. Cell Cycle 2016; 15: 1667–1673.

17. Wang Y, Yella J, Chen J, McCormack FX, Madala SK, Jegga AG. Unsupervised gene expression analysis identify IPF-severity correlated signatures, associated genes and biomarkers. BMC Pulm Med 2017; 17: 133.

18. Mullenbrock S, Liu F, Szak S, Hronowski X, Gao B, Juhasz P, Sun C, Liu M, McLaughlin H, Xiao Q, Feghali-Bostwick C, Zheng TS. Systems Analysis of Transcriptomic and Proteomic Profiles Identifies Novel Regulation of Fibrotic Programs by miRNAs in Pulmonary Fibrosis Fibroblasts. Genes (Basel) 2018; 9.

19. Hoff CR, Perkins DR, Davidson JM. Elastin gene expression is upregulated during pulmonary fibrosis. Connect Tissue Res 1999; 40: 145–153.

20. Chen W, Yan X, Xu A, Sun Y, Wang B, Huang T, Wang H, Cong M, Wang P, Yang A, Jia J, You H. Dynamics of elastin in liver fibrosis: Accumulates late during progression and degrades slowly in regression. J Cell Physiol 2019; 234: 22613–22622.

21. Singh A, Koduru B, Carlisle C, Akhter H, Liu RM, Schroder K, Brandes RP, Ojcius DM. NADPH oxidase 4 modulates hepatic responses to lipopolysaccharide mediated by Toll-like receptor-4. Sci Rep 2017; 7: 14346.

22. Amara N, Goven D, Prost F, Muloway R, Crestani B, Boczkowski J. NOX4/NADPH oxidase expression is increased in pulmonary fibroblasts from patients with idiopathic pulmonary fibrosis and mediates TGFbeta1-induced fibroblast differentiation into myofibroblasts. Thorax 2010; 65: 733–738.

23. Kakino S, Ohki T, Nakayama H, Yuan X, Otabe S, Hashinaga T, Wada N, Kurita Y, Tanaka K, Hara K, Soejima E, Tajiri Y, Yamada K. Pivotal Role of TNF-α in the Development and Progression of Nonalcoholic Fatty Liver Disease in a Murine Model. Horm Metab Res 2018; 50: 80–87.

24. Mukhopadhyay S, Hoidal JR, Mukherjee TK. Role of TNFalpha in pulmonary pathophysiology. Respir Res 2006; 7: 125.

25. Grant R, Nguyen KY, Ravussin A, Albarado D, Youm YH, Dixit VD. Inactivation of C/ebp homologous protein-driven immune-metabolic interactions exacerbate obesity and adipose tissue leukocytosis. J Biol Chem 2014; 289: 14045–14055.

26. Sun B, Wang X, Ji Z, Wang M, Liao YP, Chang CH, Li R, Zhang H, Nel AE, Xia T. NADPH Oxidase-Dependent NLRP3 Inflammasome Activation and its Important Role in Lung Fibrosis by Multiwalled Carbon Nanotubes. Small 2015; 11: 2087–2097.

27. Zhang LL, Huang S, Ma XX, Zhang WY, Wang D, Jin SY, Zhang YP, Li Y, Li X. Angiotensin(1-7) attenuated Angiotensin II-induced hepatocyte EMT by inhibiting NOX-derived H2O2-activated NLRP3 inflammasome/IL-1β/Smad circuit. Free Radic Biol Med 2016; 97: 531–543.

28. Owona BA, Abia WA, Moundipa PF. Natural compounds flavonoids as modulators of inflammasomes in chronic diseases. Int Immunopharmacol 2020; 84: 106498.

29. Kim HS, Yoon YM, Meang MK, Park YE, Lee JY, Lee TH, Lee JE, Kim IH, Youn BS. Reversion of in vivo fibrogenesis by novel chromone scaffolds. EBioMedicine 2019; 39: 484–496.

30. Nonnenmacher Y, Hiller K. Biochemistry of proinflammatory macrophage activation. Cell Mol Life Sci 2018; 75: 2093–2109.

31. Redente EF, Keith RC, Janssen W, Henson PM, Ortiz LA, Downey GP, Bratton DL, Riches DW. Tumor necrosis factor-α accelerates the resolution of established pulmonary fibrosis in mice by targeting profibrotic lung macrophages. Am J Respir Cell Mol Biol 2014; 50: 825–837.

