## supplementary figures for "A noble TGFβ biogenesis inhibitor exhibits both potent anti-fibrotic and anti-inflammatory capabilities"

## A)

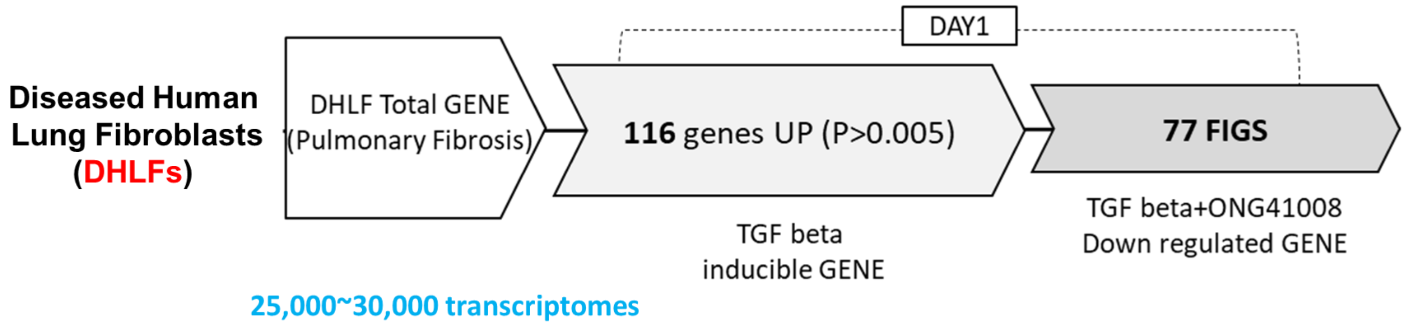


**Fibrosis-Inducing Genes (FIGS)**

#### B) Human DHLFs regulated by ONG41008 Via RNA-Seq / p< 0.005

| | |
|---|---|
| EPHB2 | PAWR |
| LEPRE1 | APCDD1L |
| KIF26B | GALNT10 |
| PLXDC2 | PKP1 |
| ANKRD1 | PXDC1 |
| POSTN | GJA1 |
| XYLT1 | NTM |
| FAM101B | SH3PXD2A |
| CDH2 | LEPREL4 |
| PXDN | PODXL |
| COL7A1 | PSAT1 |
| PCDH10 | NOX4 |
| CDH6 | RP11-180C1.1 |
| SPOCK1 | FBLN5 |
| EDN1 | ADAM19 |
| CTGF | MRC2 |
| IGFBP3 | COL1A1 |
| LRRC17 | WWTR1 |
| PLAT | LOX |
| CTHRC1 | ITGB5 |
| COL5A1 | HHAT |
| NEGR1 | KSR1 |
| VDR | HMCN1 |
| COL4A1 | GATA6 |
| TPM1 | CDK6 |
| COL11A1 | P4HA2 |
| FAM46A | EFEMP1 |
| EDIL3 | ALDH1L2 |
| ELN | SLC38A5 |
| DKK1 | SEMA7A |
| TCF4 | FIBCD1 |
| PLOD2 | NCOA3 |
| NEK7 | P4HA1 |
| | ATP10A |
| | PYCR1 |
| | RNF152 |
| | HAPLN1 |
| | COL5A2 |
| | DGKI |
| | HYOU1 |
| | ASXL3 |
| | SERPINH1 |
| | SOX2-OT |

“NOX4”

77 FIGS

**Supplementary Figure 1**

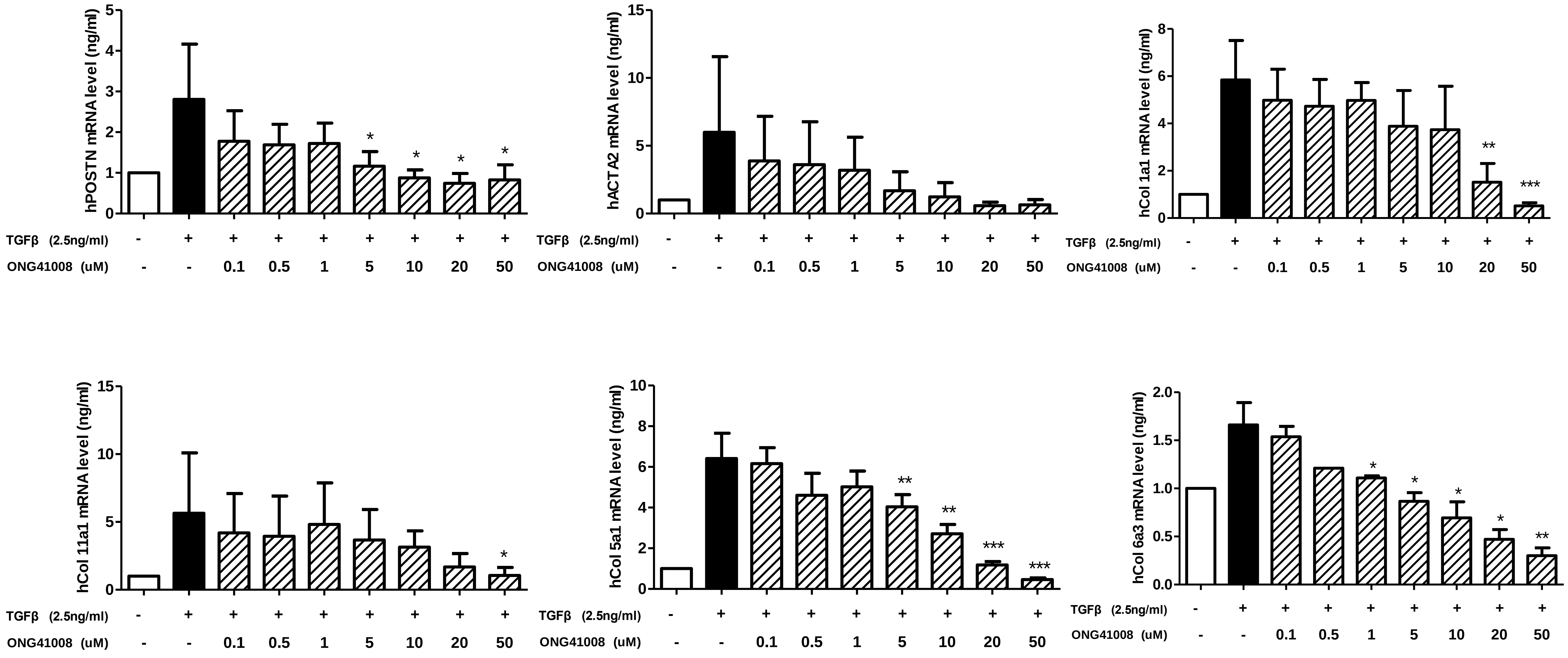


### Supplementary Figure 2

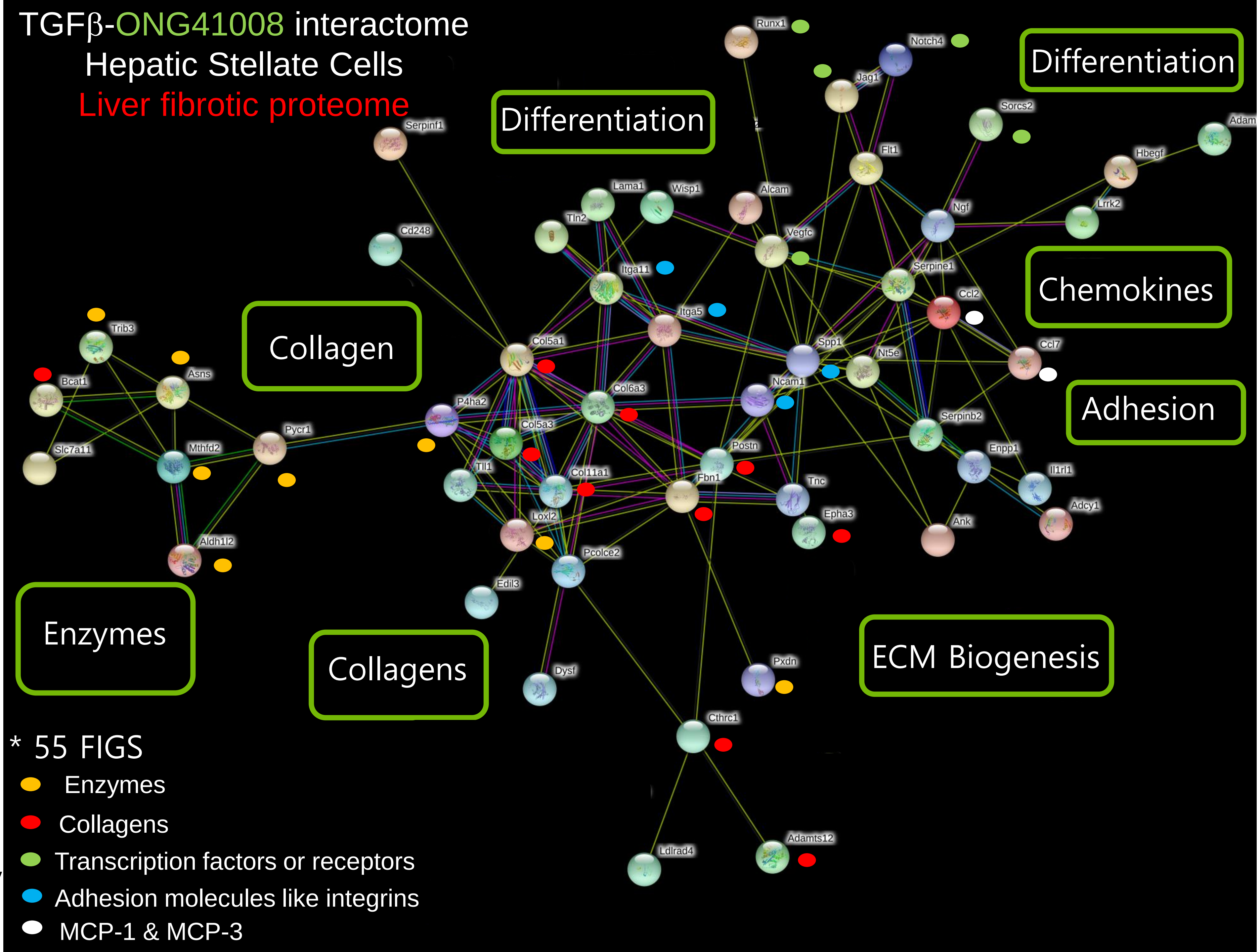


**Supplementary Figure 3**
